# N-acetylcysteine Microparticles Reduce Cisplatin-induced RSC96 Schwann Cell Toxicity

**DOI:** 10.1101/2023.10.31.564430

**Authors:** Katherine Kedeshian, Michelle Hong, Larry Hoffman, Ashley Kita

## Abstract

**Objectives:** Cisplatin is known to cause inner ear dysfunction. There is growing evidence that cisplatin-induced demyelination of spiral or Scarpa’s ganglion neurons may play an additional role in drug-induced ototoxicity alongside afferent neuron injury. As Schwann cells produce myelin, there may be an opportunity to reduce ototoxic inner ear damage by promoting Schwann cell viability. This work describes a cellular model of cisplatin-induced Schwann cell injury and investigates the ability of the antioxidant N-acetylcysteine to promote Schwann cell viability. A local delivery system of drug-eluting microparticles was then fabricated, characterized, and investigated for bioactivity.

**Methods:** RSC96 rat Schwann cells were dosed with varying concentrations of cisplatin to obtain a dose curve and identify the lethal concentration of 50% of the cells (LC_50_). In subsequent experiments, RSC96 cells were co-treated with cisplatin and both resuspended or eluted N-acetylcysteine. Cell viability was assessed with the CCK8 assay.

**Results:** The LC_50_ dose of cisplatin was determined to be 3.76 μM (p=2.2 × 10^−16^). When co-dosed with cisplatin and therapeutic concentration of resuspended or eluted N-acetylcysteine, Schwann cells had an increased viability compared to cells dosed with cisplatin alone.

**Conclusion:** RSC96 Schwann cell injury following cisplatin insult is characterized in this in vitro model. Cisplatin caused injury at physiologic concentrations and N-acetylcysteine improved cell viability and mitigated this injury. N-acetylcysteine was packaged into microparticles and eluted N-acetylcysteine retained its ability to increase cell viability, thus demonstrating promise as a therapeutic to offset cisplatin-induced ototoxicity.

**Lay Summary:** Cisplatin is a chemotherapeutic agent known to cause balance and hearing problems through damage to the inner ear. This project explored cisplatin injury in a Schwann cell culture model and packaged an antioxidant into microparticles suitable for future drug delivery applications.

## Introduction

Cisplatin is a chemotherapeutic medication known to cause ototoxicity.^1,2^ The resulting balance and hearing dysfunction cause significant disruptions to patients’ lives.^3^ Cisplatin has been shown to induce apoptosis in Schwann cells, raising the possibility that Schwann cell injury is one of the mechanisms by which cisplatin induces ototoxicity. ^4,5^ Additionally, Schwann cell damage has been shown to mimic a certain form of hearing loss known as hidden hearing loss. ^6,7^

Schwann cells make up the myelin sheath that wraps around the axon of neurons to promote nerve conduction.^8^ When damage to neurons within the peripheral nervous system occurs, free radicals are created which adversely impact Schwann cell health by inducing oxidative stress.^4^ RSC96 cells, an immortalized rat Schwann cell line, were chosen as a Schwann cell model as they have been used in other studies with cisplatin.^10^ Other studies have investigated antioxidants that blunt the effects of cisplatin ototoxicity. ^11,12^ Thus, an antioxidant like N-acetylcysteine (NAC) was hypothesized to counteract the influence of oxidative stressors. ^13–15^ A model of Schwann cell injury was then developed to investigate the protective effects of this antioxidant therapeutic.

Since a course of cisplatin therapy often requires intermittent infusions over weeks to months, one strategy to reduce cisplatin-induced ototoxicity is to design a drug-eluting microparticle able to locally deliver an otoprotective agent for a prolonged period. NAC, a hydrophilic antioxidant, was blended into poly lactic-co-glycolic acid (PLGA) microparticles. NAC was also incorporated into polycaprolactone (PCL) microparticles using a double emulsion method as previously described.^15^ Both polymers were chosen for their widely accepted use as FDA approved biocompatible polymers, making them ideal vehicles to prolong the release of a therapeutic agent able to protect patients from cisplatin-induced inner ear injury.

## Materials and Methods

### RSC96 Cell Culture

RSC96 rat Schwann cells were cultured in media consisting of Dulbecco’s modified Eagle’s medium, 1% penicillin-streptomycin and 10% fetal bovine serum. RSC96 cultures were maintained in an incubator kept at 37°C and 5% CO_2_. The reagents used are listed in Table 1.

**Table 1:**
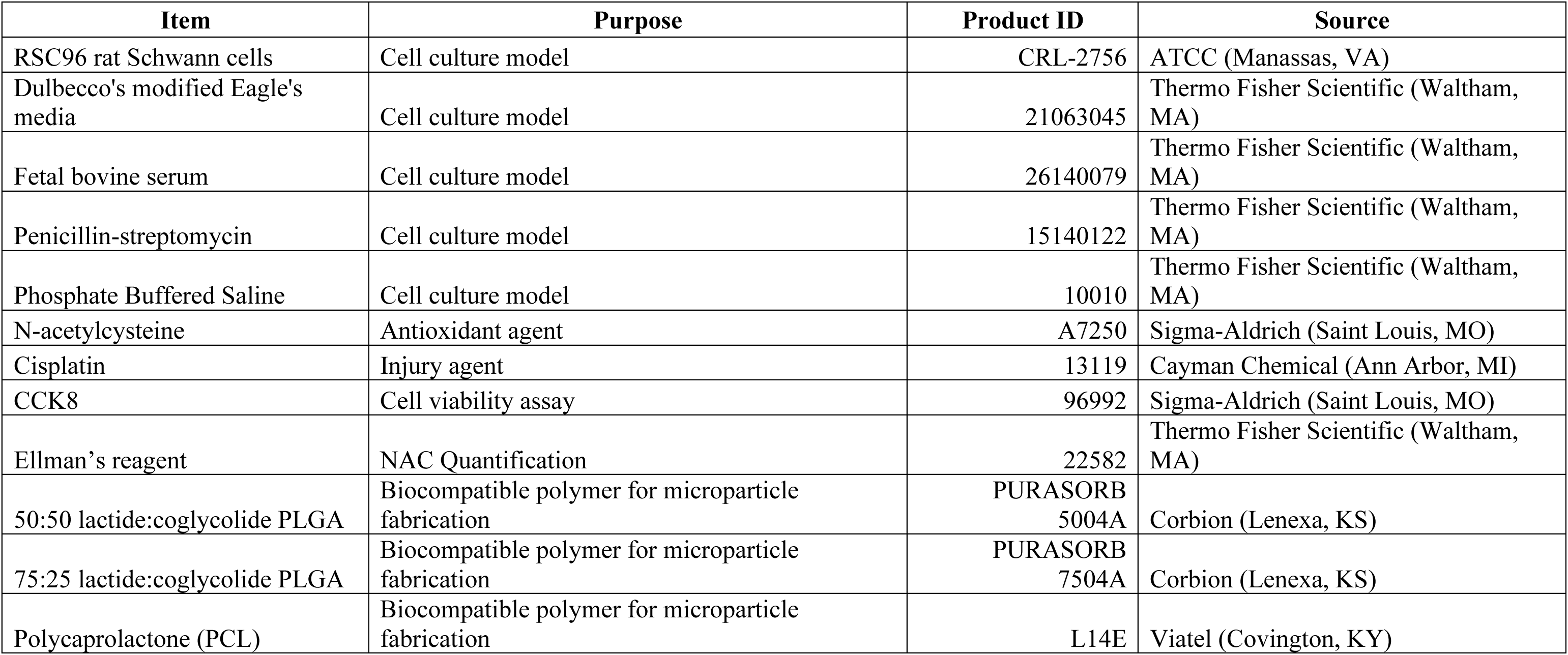
Materials Table.

### CCK8 Viability Assay

The Cell Counting Kit 8 (CCK8) assay was used to assess cell viability. This kit was used to quantify cell viability by producing a formazan dye after reduction of a tetrazolium salt.

### LC_50_ Determination

RSC96 cells were plated at a density of 4000 cells/well in a 96-well plate and were allowed to grow for 24 hours. Following the 24-hour incubation period, the media was removed, after which each well received fresh media along with doses of cisplatin ranging from 0 µM-500 µM. Cell viability was determined after 48 hours using the CCK8 assay. The LC_50_ was determined using a four-parameter logistic regression.

### NAC Screening with CCK8 Assay

Following the determination of the concentration of cisplatin resulting in 50% lethality of RSC96 cells (LC_50_), the protective capacity of NAC was assessed. RSC96 cells were plated at a density of 4000 cells/well in 96-well plates and were allowed to grow for 24 hours. Cells were then co-dosed with the cisplatin LC_50_ and increasing concentrations of NAC to obtain a total volume of 100 µL/well. After 48 hours, cell viability was determined. Six conditions were explored, each with 8 replicates: control, cisplatin alone, cisplatin with 1µM NAC, cisplatin with 10µM NAC, cisplatin with 100µM NAC, and cisplatin with 1000µM NAC.

### Microparticle Fabrication and Elution

A microparticle protocol from Murphy et al. was adapted to create PLGA and PCL microparticles as well as NAC-loaded PLGA and PCL microparticles.^15,16^ Unloaded microparticles were fabricated as controls. Three separate batches were created by dissolving the following polymer formulations in dichloromethane (DCM): 2% 50:50 lactide:coglycolide poly(lactic-co-glycolic acid) (PLGA), 2% 75:25 lactide:coglycolide PLGA, or 2% polycaprolactone (PCL). Each polymer-DCM solution was then added dropwise to a 1% solution of poly(vinyl alcohol) (PVA) and homogenized at 3000 rpm for 15 minutes. The mixture was next poured into 0.3% PVA and stirred at 700 rpm for 2 hours at 40°C. The subsequent solution was then centrifuged and washed in double-distilled water three times before resuspending the microparticle solution and freezing it overnight at −80°C. Frozen microparticles were then lyophilized and stored at −20°C.

Microparticles fabricated with NAC were created by dissolving 2% 50:50 PLGA in DCM followed by 10 mg NAC in 100 µL water. The solution was then sonicated. This was then added dropwise to a 1% solution of PVA saturated with 2.5% NAC and homogenized at 3000 rpm for 15 minutes. The mixture was next poured into 0.3% PVA with 5% NAC and stirred at 700 rpm for 2 hours at 40°C. The mixture was then centrifuged and washed in double-distilled water with 2.5% NAC three times before freezing and lyophilization. Microparticles were similarly made with 75:25 PLGA and PCL.

Approximately 10 mg of each microparticle type was suspended in 150 µL double distilled water, a sample of which was placed on a glass slide for microparticle imaging using a Zeiss AxioPlan microscope with a 20x objective. Six images of each condition were obtained to ensure broad sampling. The *imfindcircles* function (Matlab Image Processing Toolbox) was used to identify and segment the microparticles, from which radii were determined. PCL images were converted to binary arrays using Fiji’s built-in “Make Binary” function and then “Fill Holes” to prevent detection of within particle features otherwise identified as individual microparticles by the *imfindcircles* function. These within microparticle features were not detected in the PLGA microparticle images.

### Microparticle Elution and RSC96 Dosing

Lyophilized microparticles were weighed directly into 12 mm 0.4 µm-pore polyester transwell inserts. The transwell inserts were filled with 0.6 mL of phosphate buffered saline (PBS) and placed into 1.5 mL of PBS within 12 well plates. Two replicates of each PLGA microparticle condition and four replicates of each PCL condition were used to characterize elution. Eluted solution was collected by exchanging the 1.5 mL of PBS below the transwell containing microparticles and placing this within cryotubes stored at −20°C. This was done daily for the first 5 days then twice weekly for the first month and weekly thereafter. The concentration of NAC in the eluted solution was determined via linear regression following incubation with Ellman’s reagent, a compound that reacts with sulfhydryl groups to induce a color change that is then read on a plate reader at 412 nm.^15^

RSC96 cells were plated in 96 well plates at a density of 4000 cells/well for 24 hours and then co-dosed with cisplatin and resuspended NAC or eluted NAC solutions from 50:50 PLGA and PCL microparticles. As NAC was eluted into PBS, eluted solutions were diluted with media to the appropriate concentration for dosing. The CCK-8 assay was then used to determine cell viability.

### Statistical Analyses

Preliminary statistical analyses were conducted using the R Studio statistical software platform (R Core Team, 2017). A bootstrap implementation of one- and two-way analyses of variance (ANOVA) were conducted using custom scripts written in the *Igor Pro* software environment. RSC96 viability under conditions of varying NAC concentration was analyzed using a single factor ANOVA with the Dunnet’s post-hoc test. A two-way ANOVA was used to evaluate the effects of NAC source (i.e. resuspended or eluted) and concentration (0, 100, or 1000 µM; two-factor ANOVA), and Dunnet’s post-hoc analysis was used to distinguish levels of NAC concentration. Figures were created with Igor Pro (WaveMetrics, 2022) and Adobe Illustrator (Adobe Inc, 2019) and Adobe Photoshop (Adobe Inc, 2019).

## Results

### LC_50_ Determination

The LC_50_ of two experiments after normalization was 3.76 µM (Figure 1). Therefore, all subsequent experiments were conducted using the rounded value of 4 µM cisplatin.

**Figure 1.**
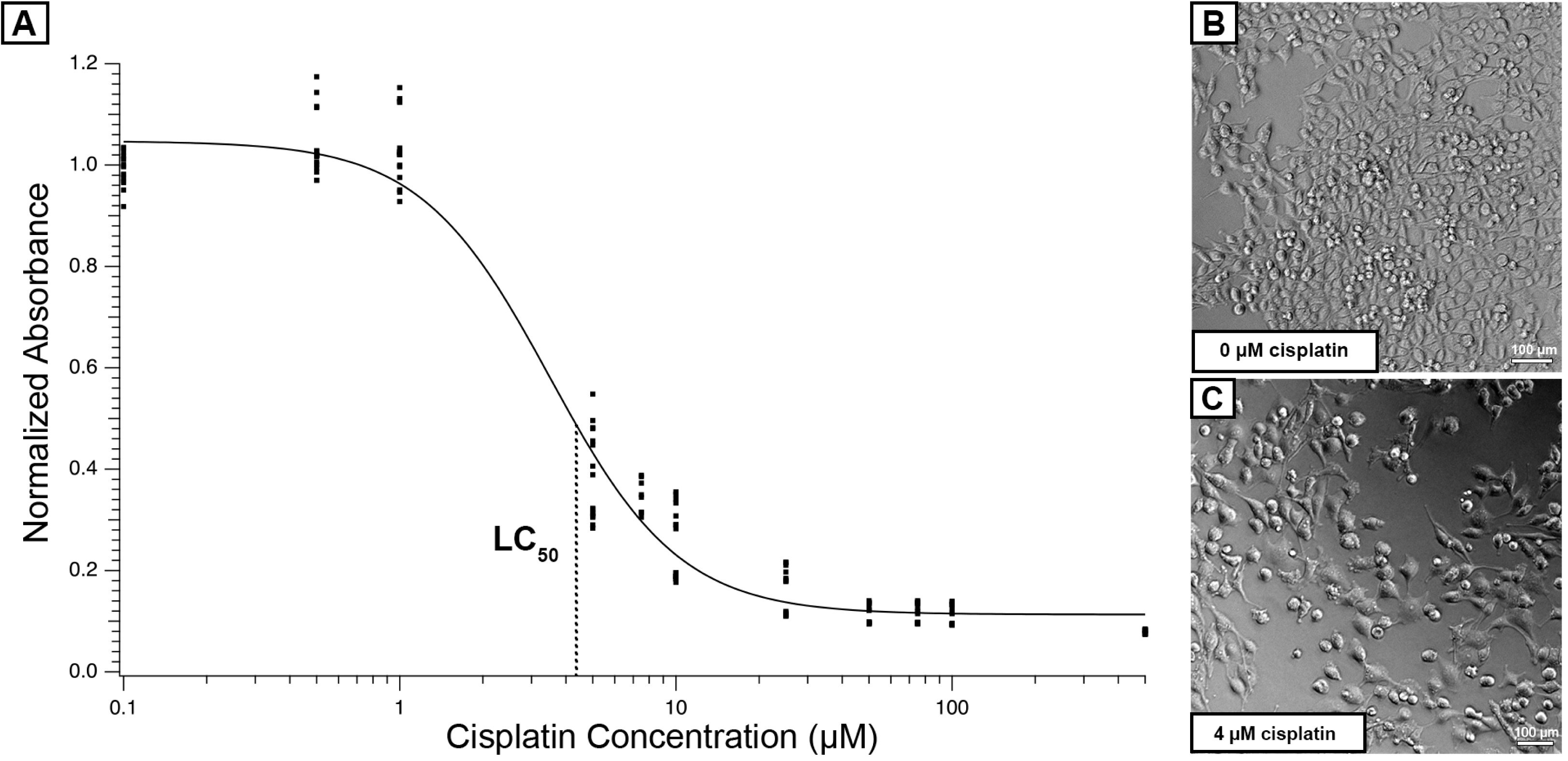
Determination of cisplatin LC_50_. A: RSC96 Cell viability at cisplatin concentrations from 1-500 µM as determined by CCK-8 assay. Curve fitted using a four-parameter logistic regression. The dose required to reduce absorbance by 50% was 3.76 µM and is indicated by a dashed line and LC_50_ label. B and C: 10X light micrographs of RSC96 cells exposed to 0 (*Control,* B) and the rounded LC_50_ value *(Cisplatin*, C).

### NAC Screening

Cells co-dosed with 100 and 1000 µM NAC with the cisplatin LC_50_ resulted in an increase in CCK8 absorbance when compared to conditions dosed with cisplatin alone (Figure 2). Cell viability was improved at NAC concentrations of 100 µM (p < 2 × 10^−6^) and 1000 µM (p < 2 × 10^−6^) compared to cells dosed with cisplatin alone (resampled single factor ANOVA with Dunnet’s post hoc test). RSC96 viability associated with NAC concentrations less than 100 µM were similar to the control condition (4 µM cisplatin and 0 µM NAC).

**Figure 2.**
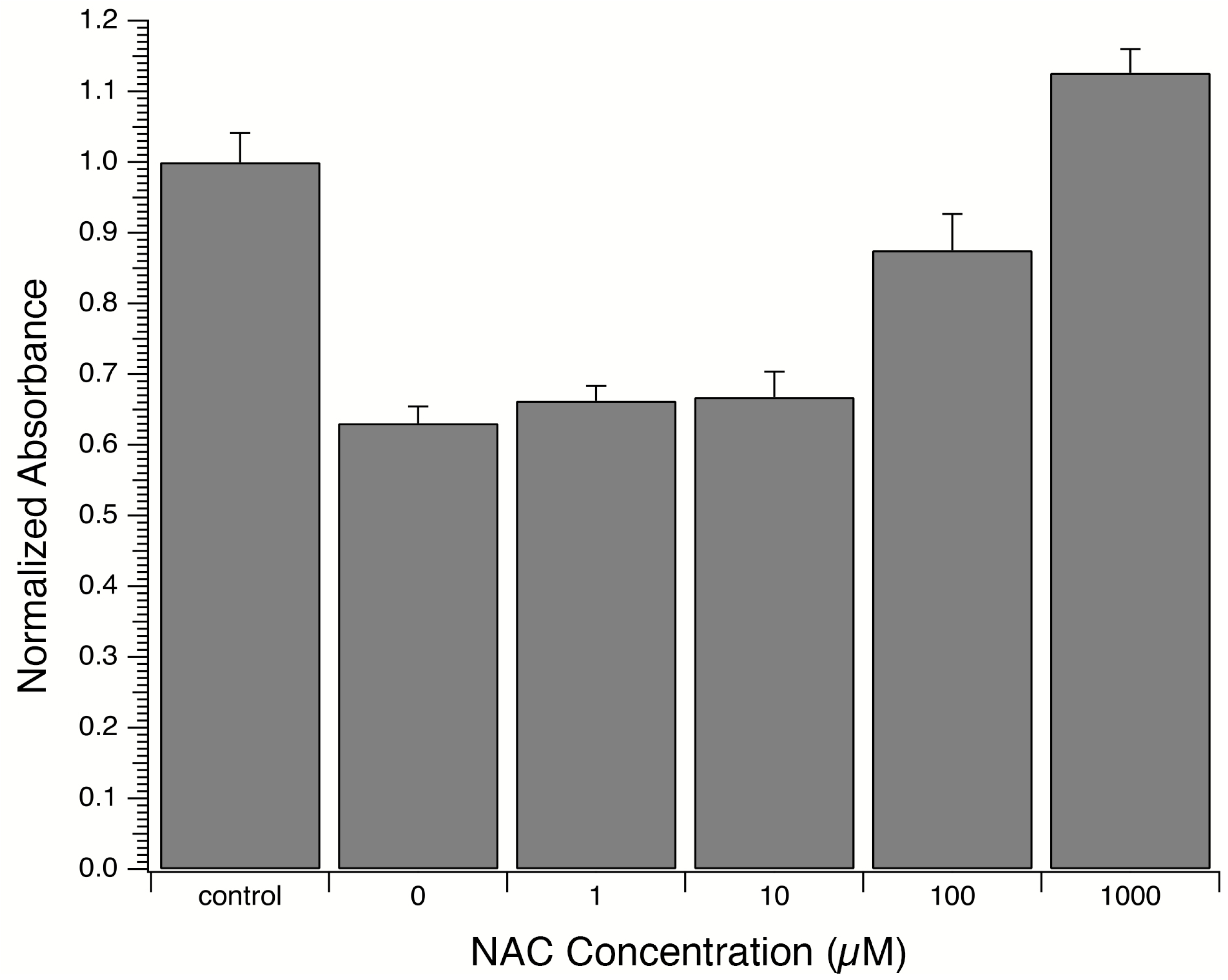
NAC Dose Screening. RSC96 cells were co-dosed with the initial LC_50_ of 3.34 μM cisplatin and different concentrations of N-acetylcysteine (NAC) (0 µM, 1 µM, 10 µM, 100 µM, 1000µM). Cells had improved viability when compared to cisplatin alone conditions at 100 µM and 1000 µM NAC.

### Microparticle Characterization

The diameters of fabricated microparticles are represented in a histogram in Figure 3.

**Figure 3.**
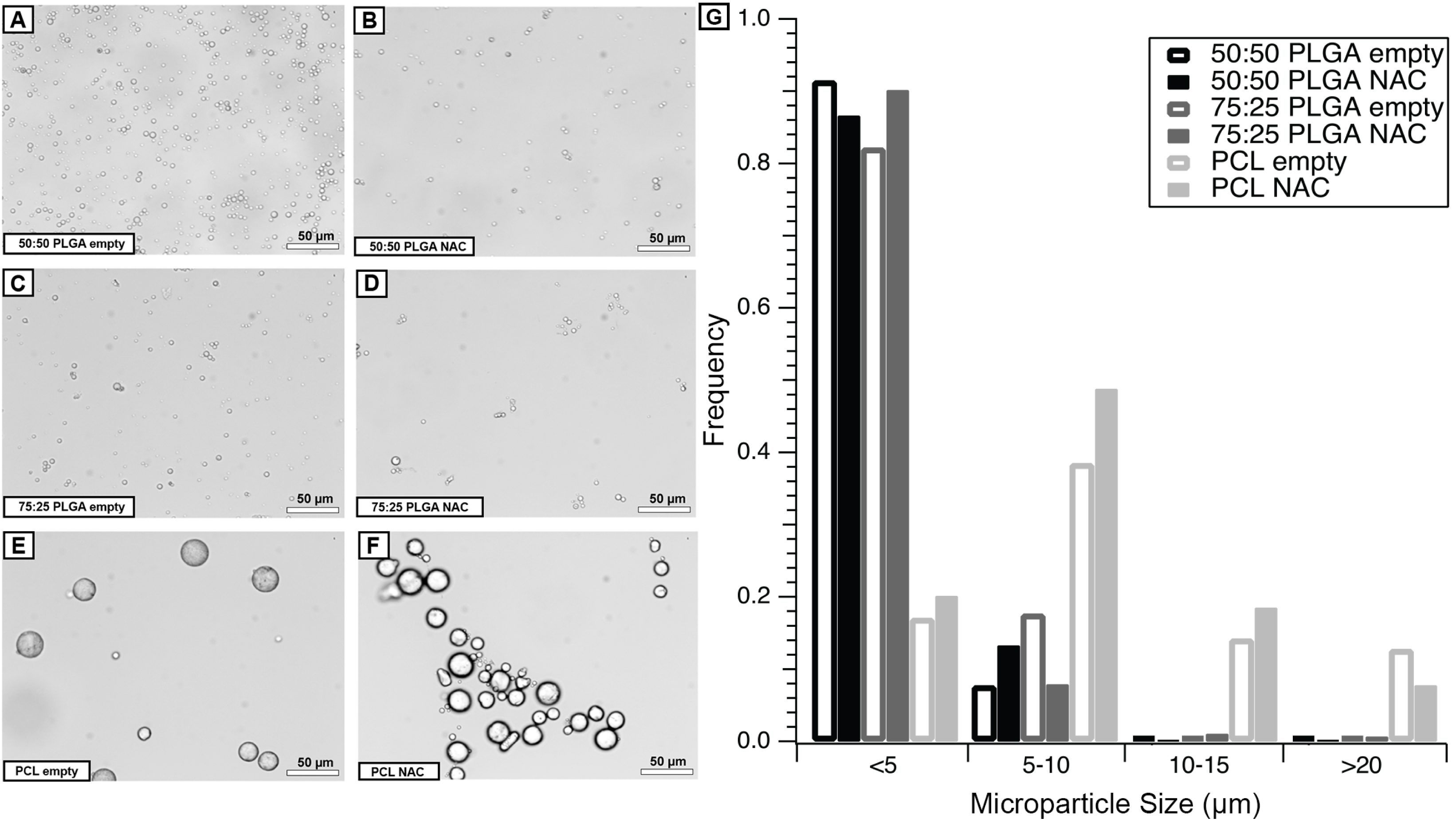
Characterization of microparticles. A-F: 20x microparticle images of each condition. G: Microparticle diameter size distribution of six 20x images per microparticle.

### Microparticle Elution

To account for the different masses of microparticles weighed into each transwell at the beginning of elution, elution is expressed as a fraction of the total amount released by the end of collection versus time (Figure 4). By three days the PLGA microparticles had all released greater than 95% of the total amount eluted. The PCL microparticle elution was slightly more prolonged releasing 83.3% of what would be released after 3 days.

**Figure 4.**
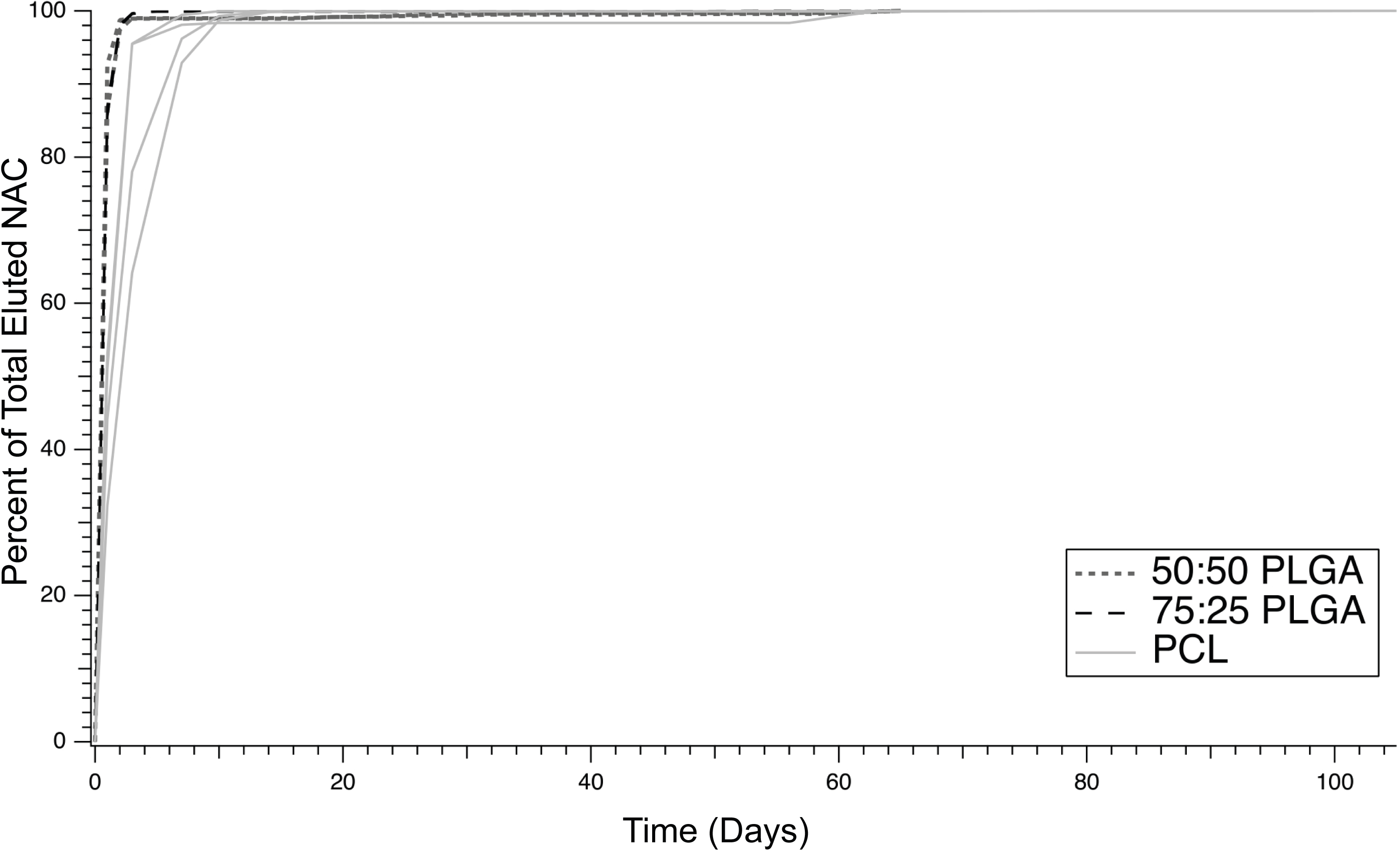
Microparticle elution. NAC elution from microparticles as a function of the total amount eluted by the end of collection to account for differences in the total amount of mass from which NAC was eluted in each transwell.

PCL microparticles with NAC incorporated released the largest amount of NAC relative to the initial mass added to their transwell, while 50:50 PLGA microparticles released the smallest amount of NAC relative to their initial mass (Figure 5).

**Figure 5.**
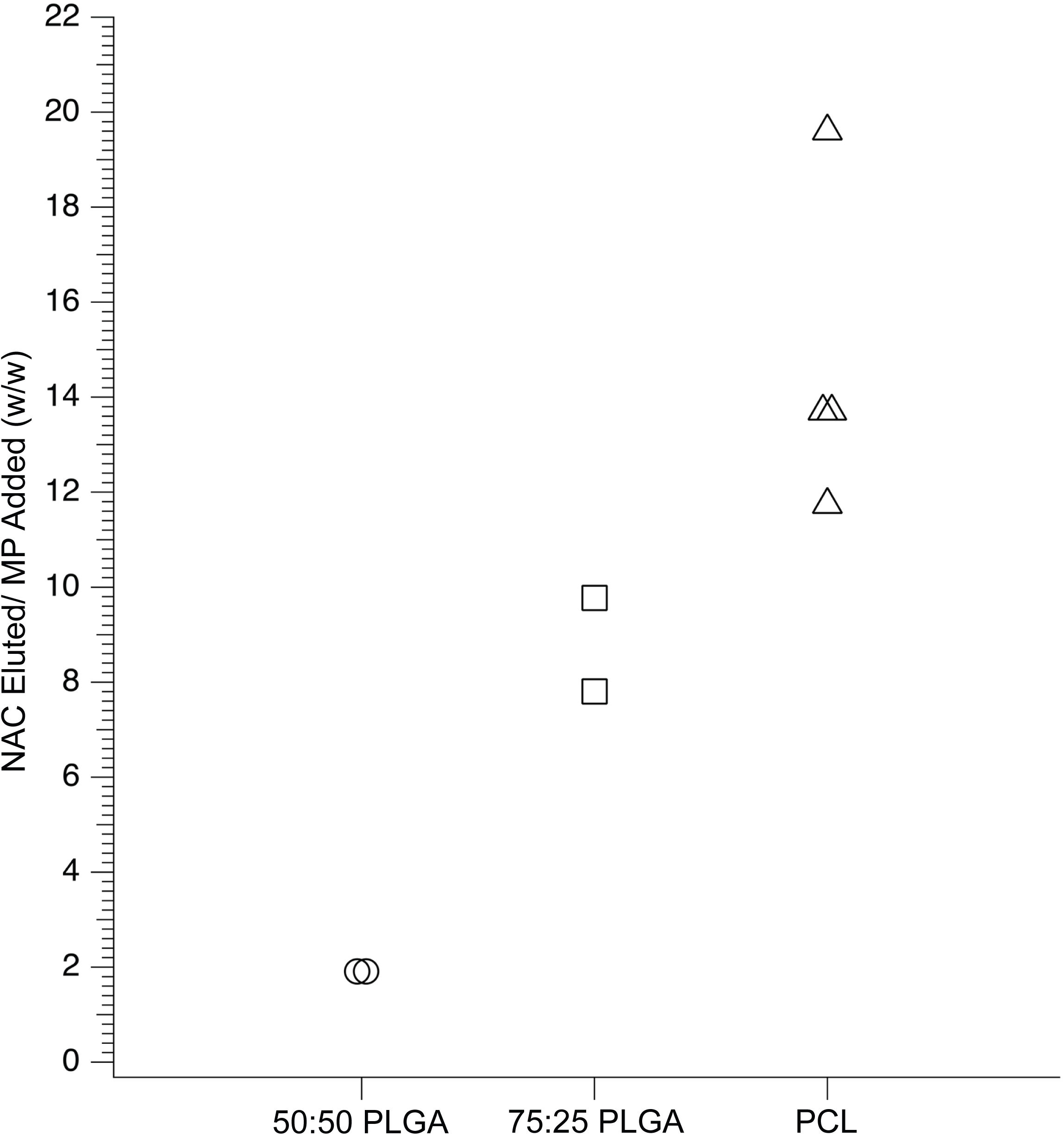
Microparticle encapsulation. Total amount of NAC released based on the total mass of microparticle condition initially added to each transwell.

### NAC Eluted from Microparticles Showed An Increase in Cell Viability

To determine whether microparticle encapsulation altered the efficacy of NAC, RSC96 cells were dosed with cisplatin along with either resuspended NAC or NAC eluted from PLGA microparticles at concentrations of 100 µM and 1000 µM and from PCL microparticles at a concentration of 100 µM (Figure 6). Additional conditions were included that dosed cells with eluted solutions from empty microparticles to establish a control absorbance and ensure that a microparticle biproduct alone was not responsible for an increase in absorbance.

**Figure 6.**
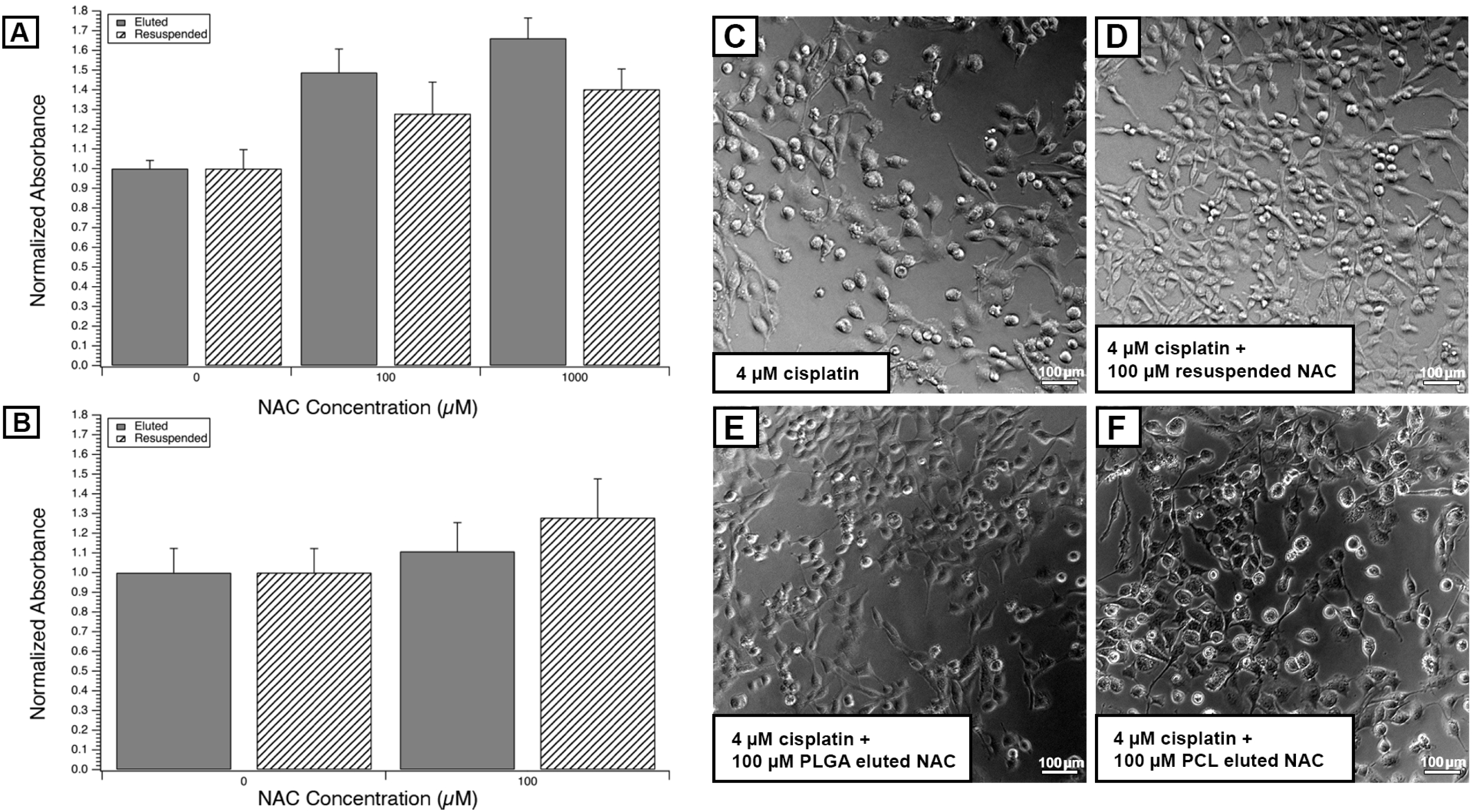
Bioactivity of eluted NAC assessment. All conditions were dosed with a rounded LC_50_ value of 4 μM of cisplatin. All replicate experiments were normalized to their respective controls. The control for eluted conditions was cells co-dosed with cisplatin and eluted solution from microparticles fabricated with no NAC. The control for resuspended conditions was cells co-dosed with cisplatin and media containing no NAC. A: NAC eluted from PLGA microparticles at 100 and 1000 µM exhibited similar increases in viability as resuspended NAC. B: NAC eluted from PCL microparticles at 100 µM exhibited similar increases in viability as resuspended NAC. C-F: Light micrographs (10X) of RSC96 cells dosed with cisplatin at 4 µM (C), cisplatin at 4 µM and resuspended NAC (D), cisplatin at 4 µM and eluted solution from PLGA microparticles at a concentration of 100 µM (E), and cisplatin at 4 µM and eluted solution from PCL microparticles at a concentration of 100 µM (F).

A protective effect was seen with NAC, whether from microparticles or resuspended. This was seen in both PLGA microparticles and PCL microparticles. Protection was seen with NAC eluted from PLGA microparticles at concentrations of 100 µM (p < 2 × 10^−6^) and 1000 µM (p < 2 × 10^−6^). NAC eluted from PCL microparticles increased viability at a concentration of 100 µM (p < 1 × 10^−4^). Additionally, there was a difference between cell viability in conditions dosed with and without eluted solutions from microparticles. Cells were unable to be co-dosed with cisplatin and 1000 µM NAC from PCL microparticles due to the low concentration of eluted solution, which would not allow for diluting and dosing cells with 1000 µM without risk of significant media deprivation.

## Discussion

### Development of a Schwann Cell Model of Cisplatin-Induced Injury

Given the deleterious effects of cisplatin on the inner ear, a cellular model can be a helpful first step in identifying therapeutic agents to mitigate this effect. In this RSC96 Schwann cell model the LC_50_ of cisplatin was 4 µM. Since standard chemotherapy treatments in humans attain a cisplatin concentration of 0.41 µM – 9.27 µM^17^, this experiment suggests that Schwann cells may undergo damage during chemotherapeutic treatment at concentrations attained therapeutically. The findings of the present study, therefore, may harbor high translational impact.

### NAC as a Therapeutic Agent

The antioxidant NAC was explored for its ability to reduce Schwann cell damage. Two investigated concentrations of NAC were found to reduce cellular injury from cisplatin. NAC also did not demonstrate toxicity in this assay at concentrations approaching its solubility limit, a desirable property when considering that prolonged local delivery often requires high initial concentrations to attain sustained therapy. Furthermore, NAC eluted from both PLGA and PCL microparticles continued to demonstrate a protective effect at the concentrations identified during the screening assay. This is encouraging that the microparticle fabrication process did not affect the bioactivity of the eluted NAC, nor did microparticle degradation during NAC elution release any byproducts that negatively impacted NAC’s viability enhancement.

### Designing a Method of Prolonging N-acetylcysteine Delivery with Microparticles

Microparticles synthesized with NAC eluted therapeutic concentrations (100 µM or greater) for 2-3 days from PLGA microparticles, and for up to 7 days from PCL microparticles. All conditions exhibited a large initial “burst release” with the most sustained release from PCL. Since variable amounts of microparticles were weighed into each condition’s transwell before measuring elution, the concentrations eluted were not directly comparable. The transwells containing PCL microparticles had a much lower microparticle mass loaded into the transwells, a result of lower overall yield in those conditions. When this is considered, it is likely that a larger mass of PCL, on the order of that weighed out for the PLGA conditions, would result in a therapeutic concentration of NAC for a period of time longer than 7 days.

The average diameter of PLGA microparticles was smaller than that of PCL. The PLGA microparticles were intentionally fabricated to be larger than those previously reported by our laboratory (3-4 µm versus 1 µm) in an effort to encapsulate more NAC and prolong elution, but this did not appear to significantly change the release profile.^15^

Since PCL microparticles had the longest sustained release, it is likely that their larger microparticle size allowed for more encapsulation of NAC and a longer duration of release as the microparticle degraded.

The duration of therapeutic concentrations of NAC delivered may be increased in the future by increasing the mass of NAC microparticles added to each transwell. Larger masses of NAC microparticles eluted into smaller volumes may allow the concentration delivered to exceed 100 µM for longer periods of time. As the volume of the perilymph in the inner ear is 158.5 µL and we eluted into solutions of 1500 µL, the same weight of microparticles could attain a concentration of 100 µM for a longer period of time in a smaller more physiologic volume.^18^

### Future Applications

For future *in vivo* applications, PCL microparticles would be more suitable. PCL containing drug delivery systems demonstrate less of an inflammatory response than PLGA microparticles.^19,20^ Additionally, PCL degrades at a slower rate in part because of its hydrophobic properties and higher molecular weight, making it more advantageous for a clinical application where chemotherapy treatment occurs over weeks to months.^21,22^ The PCL microparticles also eluted a larger amount of NAC per weight.

## Conclusion

Patients given cisplatin-based chemotherapy could benefit from the local administration of an agent that protects their inner ear from damage, ideally for a period of weeks to months. RSC96 Schwann cells were used as a model to investigate cisplatin-induced Schwann cell injury. A promising therapeutic, N-acetylcysteine, was further investigated both alone and when blended into several types of drug-eluting microparticles. The ability of this therapeutic to demonstrate protection of RSC96 cells is encouraging that this agent may be an effective method of reducing cisplatin-induced Schwann cell injury.

